# Sleep spindle deficits in childhood absence epilepsy improve with antiseizure treatment and disease resolution

**DOI:** 10.64898/2026.09.17.752393

**Authors:** Skyler K. Goodman, Sarah Yi, Dhinakaran M. Chinappen, Mark A. Kramer, Catherine J. Chu, Hunki Kwon

## Abstract

Childhood absence epilepsy (CAE), the most common childhood epilepsy syndrome, is associated with bursts of generalized thalamocortical spike-wave discharges and cognitive impairment, yet the physiological mechanisms linking disease activity, treatment, and cognition remain unclear. Sleep spindles, generated by thalamocortical circuits, are associated with cognitive function and may be disrupted by epileptic activity. Here, we retrospectively studied 87 EEGs from 53 children with active and resolved CAE and 87 age- and sex-matched controls to evaluate sleep spindle activity across disease states and its association with antiseizure medication (ASM) exposure. We found that children with active CAE had reduced spindle rates across all cortical regions compared to those with resolved CAE and controls, with the largest reduction in the frontal region, where epileptic spike activity was also most prominent. Spindle rate was inversely correlated with epileptic spike rate, consistent with a shared thalamocortical circuitry. The association between ASM exposure and spindle rate varied by disease state. In active CAE, treated children had higher spindle rates than untreated children. In resolved CAE, treated children had lower spindle rates than untreated children. In contrast, ASM-related differences in spike rate were more limited, although a medication-specific reduction was observed in active CAE. These findings support sleep spindle rate as a candidate physiological marker of disease state and ASM exposure in children with CAE. Given the established role of sleep spindles in memory consolidation, these findings support testing of whether spindle disruption provides a mechanistic link between epileptic thalamocortical activity and cognitive vulnerability in CAE.

**Significance Statement:** Childhood absence epilepsy (CAE) is a common childhood generalized epilepsy characterized by abnormal thalamocortical activity and cognitive difficulties. Here, we show that sleep spindle activity is globally reduced during active CAE, particularly in frontal regions, and normalizes after disease resolution. Spindle activity was anticorrelated with epileptic spike rate and impacted by antiseizure medication exposure. These results support the hypothesis that epileptic spikes and sleep spindles share thalamocortical circuitry and promote sleep spindles as a candidate physiological biomarker to link epileptic activity with cognitive vulnerability in CAE.

## 1. Introduction

Childhood absence epilepsy (CAE) is a subsyndrome of idiopathic generalized epilepsy and the most common pediatric epilepsy syndrome, accounting for approximately 18% of children with epilepsy^1, 2^. CAE is a self-limited disease that primarily affects school-age children 4–13 years of age^1, 3^, and is defined by frequent daily absence seizures characterized by unresponsiveness and loss of awareness accompanied by bifrontally predominant ∼3 Hz generalized spike-and-wave discharges on EEG^1, 4-7^ that cause brief behavioral arrests with loss of consciousness^5, 8^. Approximately 60% of these otherwise typically developing children are also affected by severe cognitive comorbidities^8, 9^, particularly impairments in attention^10, 11^ and memory^12, 13^. Although seizures in CAE are often pharmacologically responsive and may remit with age, cognitive dysfunction is a frequent and clinically significant comorbidity, with consistent deficits in attention and executive function observed even prior to treatment^10, 11, 14^. Further, attention deficits often remain despite seizure freedom^10, 15^. Importantly, these cognitive impairments may persist despite adequate seizure control, indicating that seizure suppression alone is insufficient to normalize cognitive outcomes^10^.

Sleep spindles are prominent 0.5-3 s sigma band (10-15 Hz) oscillatory bursts^16-19^ observed in EEG recordings during stage 2 non-rapid eye movement (NREM) sleep. Spindles are generated by well-characterized thalamocortical circuits across cortical and subcortical regions^20-25^ and are evident by approximately 2 months of age^17^. Spindle rate and spindle coupling are associated with IQ^19, 26, 27^ and more specifically reflect sleep-dependent memory consolidation across both procedural and declarative domains^27-30^. Because epileptic spikes arise within this same thalamocortical circuitry, recent work suggests that epileptic spikes can hijack the mechanisms that normally generate sleep spindles in animal models^31-34^ and in epilepsy^26, 30, 35-38^. Importantly, reduced spindle rates have been consistently associated with cognitive dysfunction^26, 36, 39^ and impaired sleep-dependent memory consolidation^30^ in children with epilepsy. Using a sleep spindle detector validated for pediatric populations and EEGs with epileptic spikes, we have previously shown that sleep spindle rates predict cognitive function more reliably than epileptic spike rates in related developmental epilepsies, including self-limited epilepsy with centrotemporal spikes^26, 30^ and epileptic encephalopathy with spike-wave-activation during sleep^36, 40^. In addition one study found that sleep spindles may be reduced in CAE^41^.

Antiseizure medication (ASM) is the primary treatment for children with CAE^11, 42, 43^. However, the impact of ASMs on cognition, particularly attention, is less well understood. In a randomized controlled trial evaluating valproic acid, ethosuximide, and lamotrigine in CAE, attentional dysfunction remained prevalent across treatment groups but was more frequent with valproic acid than with ethosuximide, though both medications provided similar seizure control^11^. Thus, a better understanding of the relationship between ASM exposure and cognition in children with CAE is needed.

In this study, we hypothesized that children with CAE would have reduced sleep spindle rates during the active disease state compared to after disease resolution. We further examined associations between ASM exposure and sleep spindle rate in both disease states and the relationship between spindle and epileptiform spike rates. Together, these analyses test whether sleep spindle rate may serve as a candidate physiological marker of disease state and ASM exposure in children with CAE.

## 2. Methods

### Subjects

All patients who underwent EEG recordings in the Massachusetts General Hospital (MGH) EEG Lab from March 2002 through March 2024 who were diagnosed with CAE by a pediatric neurologist, had no unrelated neurological conditions, and had available EEG recordings that captured stage 2 NREM sleep were eligible for inclusion. All available EEG recordings meeting these criteria were included, allowing individual children to contribute multiple recordings. Clinical information for each patient, including age, sex, ASM treatment, and disease stage at the time of EEG acquisition, was collected through review of the electronic medical record. Patients with neurological diagnoses unrelated to CAE, unavailable EEG recordings, or recordings without stage 2 NREM sleep were excluded from analysis. Resolved CAE was defined as seizure freedom for at least 12 months before EEG^44^, whereas active CAE was defined as the presence of at least one absence seizure within the preceding 12 months.

Control subjects were selected from a previously published pediatric EEG cohort^17^. Briefly, the cohort included children who underwent clinical EEG recordings in the MGH EEG Lab from February 2002 through June 2021 with available recordings that captured stage 2 NREM sleep and met criteria for no neuroactive medication use during the recording period and documented normal neurodevelopment from birth through follow-up. For the current study, one control participant was matched for each EEG recording from patients with active or resolved CAE based on the same sex and the nearest age at EEG acquisition, calculated in years and months. Each control participant was selected only once. When multiple eligible controls of the same sex had the same minimum age difference, one control was selected at random.

### EEG recordings

Clinical EEG recordings (Xltek, Natus Medical) were obtained using the 10-20 electrode system (Fp1, Fp2, F3, F4, C3, C4, P3, P4, F7, F8, T3, T4, T5, T6, O1, O2, Fz, Cz, and Pz). Sampling frequencies ranged from 256 to 1024 Hz. EEG electrodes were applied by MGH EEG technicians and impedances were maintained below 10 kΩ. Data analyses were conducted following protocols approved and monitored by the Massachusetts General Hospital Institutional Review Board according to National Institutes of Health guidelines.

### Automated spindle detection

To detect sleep spindles, EEG data during stage 2 NREM sleep were identified by visual inspection by a board-certified clinical neurophysiologist (CJC). Data were re-referenced to the common average. Sleep spindles were detected using an automated algorithm trained and validated on approximately 20,000 manually identified sleep spindles from 115 children with and without epilepsy^17, 26^. Briefly, EEG signals were segmented into 0.5 s epochs, from which theta-band power (4–8 Hz), sigma-band power (9–15 Hz), and the Fano factor of oscillatory cycle intervals were extracted to quantify cycle-to-cycle regularity^26^. The 0.5 s epoch length was chosen to match the minimum duration typically used to define sleep spindles and to provide a 2 Hz frequency resolution for stable estimation of theta activity^45^. Detected spindles were required to last at least 0.5 s, and events occurring within 1 s were merged and considered a single spindle. Spindle rate was calculated for each electrode and subsequently averaged across predefined regions, including frontal, central, temporal, parietal, and occipital regions^17, 26^.

### Automated spike detection

Spike detection used an automated algorithm described previously and modified to enable channel-wise detection^46^. The same patient datasets, time segments, and electrode channels used for spindle analysis were analyzed for spike detection. Spike rate was calculated for each electrode and then averaged within the same predefined cortical regions used for the spindle analysis.

### Statistical analysis

To test differences in spindle rate between groups (control, resolved CAE, and active CAE), we applied a linear mixed-effects model with spindle rate as the dependent variable and group as the main predictor, adjusting for age and including a subject-specific random intercept to account for repeated measures per subject. The model was applied to spindle rates averaged across channels within each cortical regions (frontal, central, temporal, parietal, and occipital) to explore regional variations in spindle rate across groups.

To evaluate whether spindle reductions were more prominent in frontal regions compared with other brain regions, spindle rates were standardized as z-scores relative to the control group, and frontal z-scores were compared with central, temporal, parietal, and occipital regions using paired t-tests. False discovery rate (FDR) correction was applied to account for multiple comparisons, with statistical significance defined at p < 0.05.

To test for a relationship between spindle rate and spike rate, we estimated a generalized linear mixed-effects model (binomial distribution and logit link). Spindle probability per second was the dependent variable, and spike rate was the main predictor, adjusting for age and including a subject-specific random intercept to account for repeated measures per subject. Model coefficients were exponentiated to obtain the odds ratios of spindle probability per second.

To test for differences in spike rate between active and resolved CAE, spike rates were first transformed using the inverse hyperbolic sine (IHS) to adjust for the right-skewed distribution of values and inclusion of zeros. We applied a linear mixed-effects model for each cortical region (frontal, central, temporal, parietal, and occipital) with IHS-transformed spike rate as the dependent variable and disease state as the main predictor, adjusting for age and including a subject-specific random intercept to account for repeated measures.

To evaluate whether spike rates were higher in frontal regions, IHS-transformed frontal spike rates were compared with central, temporal, parietal, and occipital spike rates using paired t-tests. FDR correction was applied to account for multiple comparisons, with statistical significance defined as an FDR-adjusted p < 0.05.

To test the association of ASM treatment with spindle rate (or IHS-transformed spike rate) in resolved or active CAE, we applied a linear mixed-effects model with spindle rate (or IHS-transformed spike rate) as the dependent variable and ASM as the main categorical predictor, adjusting for age and including a subject-specific random intercept to account for repeated measures per subject.

### Data availability

Derived data supporting the findings of this study are available from the corresponding author upon reasonable request. The code for spindle detection is available at https://github.com/Mark-Kramer/Spindle-Detector-Method and the code for spike detection is available at https://github.com/erinconrad/FC_toolbox/tree/main/spike_detector.

## 3. Results

### Subject characteristics

A total of 140 unique children were included in this study: 53 children with CAE contributed 87 EEG recordings (mean age 8.3 years, range 4.1-15.8 years, 60 female) and 87 controls (mean age 8.3 years, range 4.1-15.9 years, 60 female). Among children with CAE, 31 contributed one recording and 22 contributed multiple recordings (mean 2.6, range 2-4). Of the 87 CAE recordings, 59 were obtained during active disease and 28 during resolved disease; 52 were obtained during ASM treatment and 35 without ASM treatment. Subject characteristics are provided in **Table 1**.

**Table 1.** Subject characteristics.

|  | Active |  | Resolved |  | Control |
| --- | --- | --- | --- | --- | --- |
|  | Untreated | Treated | Untreated | Treated |  |
| **Number of EEG recordings** | 25 | 34 | 10 | 18 | 87 |
| Female sex (%) | 76.0 | 67.6 | 60.0 | 66.7 | 69.0 |
| Sleep Duration (min) | 18.9±7.2 | 18.1±7.9 | 16.7±6.8 | 18.5±7.3 | 16.5±7.7 |

### Children with active CAE have lower sleep spindle rates across cortical regions

Children with active CAE had lower sleep spindle rates than controls (p = 9.9×10^-6^, mean decrease 1.7 spindles/min; **Figure 1C**). This difference was observed in each brain region evaluated: frontal (p = 2.2×10^-6^, mean decrease 2.1 spindles/min), central (p = 2.8×10^-4^, mean decrease 1.5 spindles/min), temporal (p = 2.7×10^-5^, mean decrease 1.8 spindles/min), parietal (p = 6.9×10^-4^, mean decrease 1.4 spindles/min), and occipital (p=0.01, mean decrease 1.0 spindles/min).

**Figure 1.**
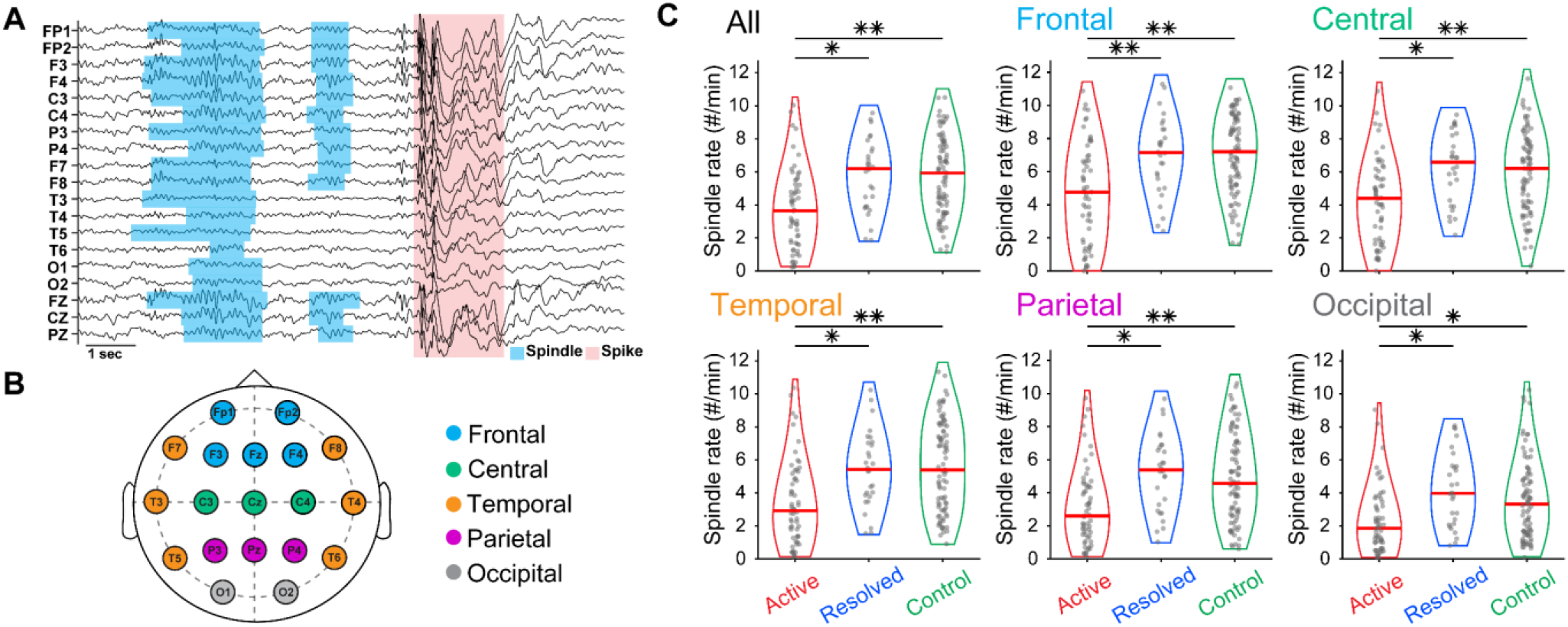
Reduced sleep spindle rate in children with active CAE. **A)** Example sleep spindles (blue shading) and generalized spike–wave discharges (red shading) recorded from an 8-year-old child with active CAE. **B)** Map of electrodes in the standard 10-20 EEG system, color-coded by cortical region; see legend. **C)** Sleep spindle rate is reduced in each cortical region in children with active CAE compared to children with resolved CAE and controls. Each violin plot shows the distribution of spindle rates for each group, with the red horizontal line indicating the median. *p<0.05, **p<0.001

In addition, children with active CAE had lower spindle rates than children with resolved CAE (p=0.002, mean decrease 1.2 spindles/min). This difference was also observed across all brain regions: frontal (p = 9.3×10^-4^, mean decrease 1.5 spindles/min), central (p=0.006, mean decrease 1.1 spindles/min), temporal (p=0.01, mean decrease 1.0 spindles/min), parietal (p=0.02, mean decrease 0.9 spindles/min), and occipital (p=0.003, mean decrease 1.2 spindles/min) regions.

### Spindle deficits are most pronounced frontally in active CAE

We found no evidence that spindle rate distributions violated the assumption of normality in any cortical region for any group (KS-test, all p > 0.17; **Figure 2A**. Spindle rates of children with active CAE normalized relative to controls were significantly lower in the frontal region than in each other region: central (t = -3.5, p = 5.2×10^-4^), temporal (t = -2.5, p = 0.007), parietal (t = -3.6, p = 3.4×10^-4^), and occipital regions (t = -5.1, p = 2.1×10^-6^; **Figure 2B**). In resolved CAE, there was a trend toward lower frontal spindle rates compared with central (t = -2.0, p = 0.03) and occipital rates (t = -2.3, p = 0.02), but neither difference was significant after FDR correction (FDR-adjusted p = 0.051 for both). Frontal spindle rates did not differ significantly from temporal (p = 0.7) or parietal rates (p = 0.1; **Figure 2C**).

**Figure 2.**
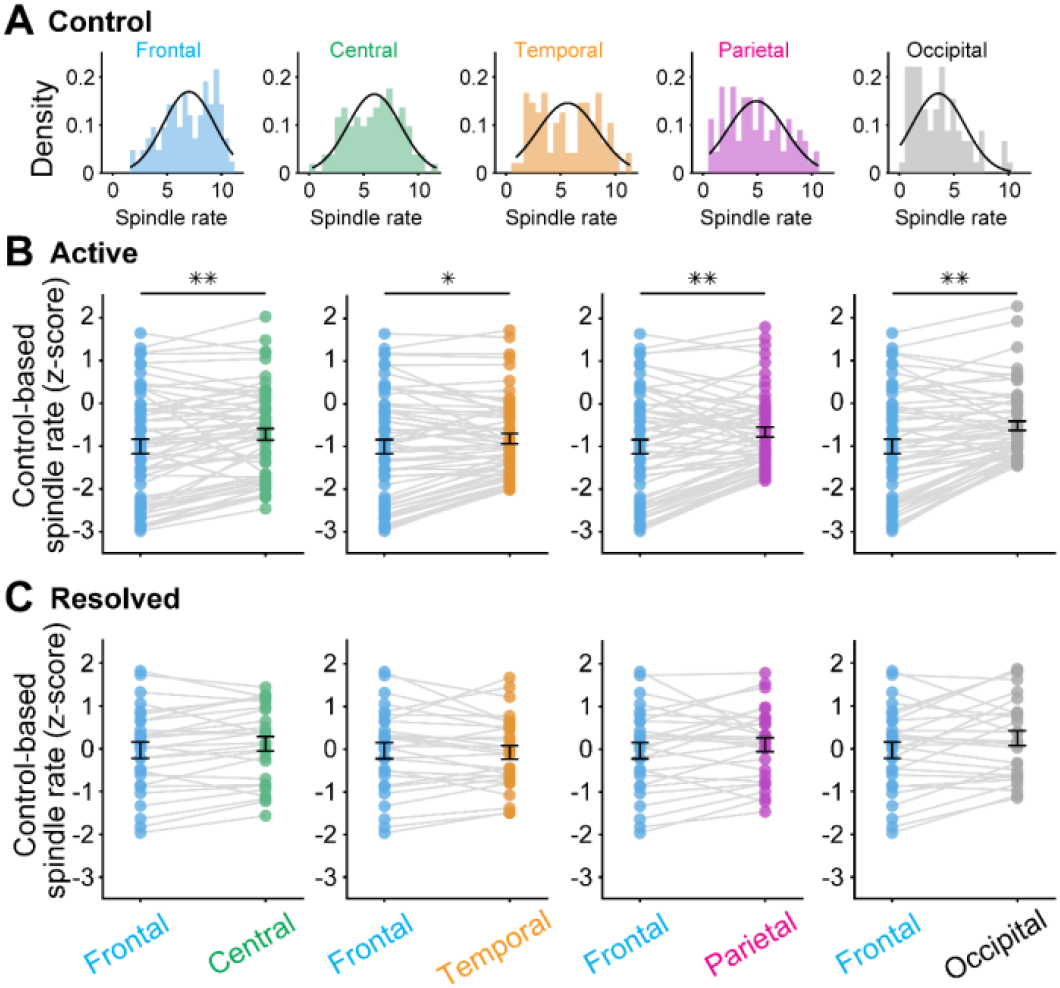
Frontal predominance of spindle deficits in active CAE. **A)** Distribution of sleep spindle rates across cortical regions in controls. (**B, C**) Spindle rates in active **(B)** and resolved **(C)** CAE were normalized to controls (z-scores). Black bars indicate mean ± SEM and gray lines indicate paired regional measurements. Frontal z-scored spindle rate was significantly lower than the z-scored spindle rates in the central, temporal, parietal, and occipital regions in active CAE, but not in resolved CAE. *p<0.05, **p<0.001

### Spike activity shows frontal predominance in active CAE and is associated with reduced spindle rate

Spike rates were significantly higher in children with active CAE than in those with resolved CAE across all cortical regions (all p < 0.001; **Figure 3A**). Within active CAE, frontal spike rates were higher than those of the central (t = 6.7, p = 5.4×10^-9^), temporal (t = 4.5, p = 1.6×10^-5^), parietal (t = 4.1, p = 7.0×10^-5^), and occipital regions (t = 2.5, p = 0.007; **Figure 3B**). In contrast, in resolved CAE, there was a trend toward higher frontal spike rates than temporal rates (p = 0.03), but this difference was not significant after FDR correction (FDR adjusted p = 0.12). Frontal spike rates did not differ significantly from central (p = 0.1), parietal (p = 0.8), or occipital rates (p = 0.06, **Figure 3C**).

**Figure 3.**
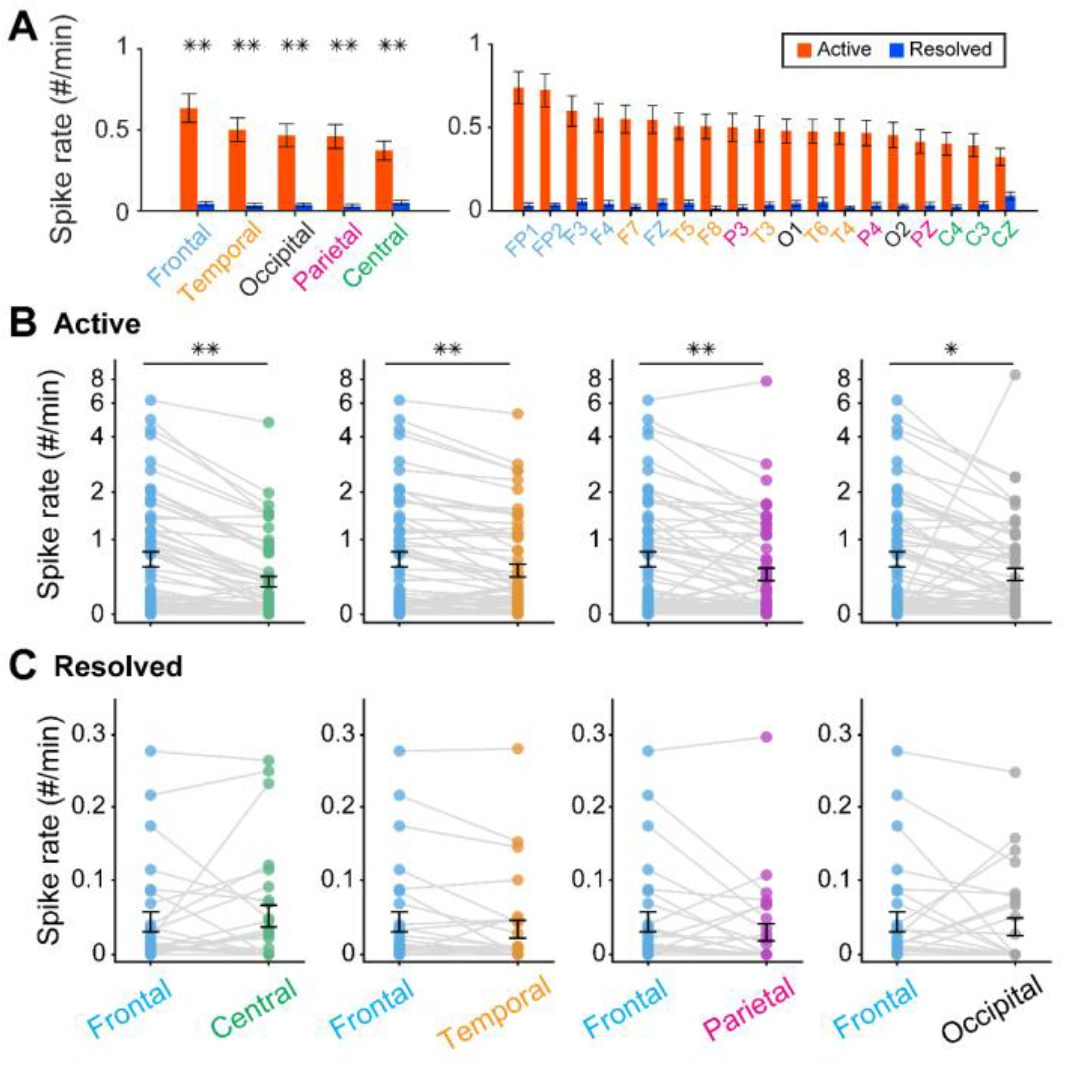
Frontal predominance of spike activity in active CAE. **(A)** Regional spike rates in children with active and resolved CAE. Lobes and electrodes are ordered by mean spike rate in active CAE, from highest to lowest. Spike rates were consistently higher in active than resolved CAE across all cortical regions. Error bars indicate mean ± SEM. **(B)** Within children with active CAE, spike rates were higher in the frontal region compared to all other regions. **(C)** Within children with resolved CAE, frontal spike rates did not differ significantly from those of other cortical regions. Each point represents an individual participant, and gray lines connect paired regional measurements within the same participant. Black bars indicate mean ± SEM. **p < 0.001.

In children with CAE, a 1 Hz increase in frontal spike rate was associated with a mean 21% reduction in the odds of spindle occurrence per second (p = 2.9×10^-4^, 95% CI [10%, 33%] reduction; **Figure 4**). A secondary analysis restricted to children with active CAE indicated a consistent – but weaker – association in the frontal region (mean 12% reduction, p = 0.051, 95% CI [0%, 23%] reduction). We conclude that higher spike rates were associated with lower odds of spindle occurrence in CAE, consistent with patterns reported across epilepsy populations^26, 30, 35-38, 40^.

**Figure 4.**
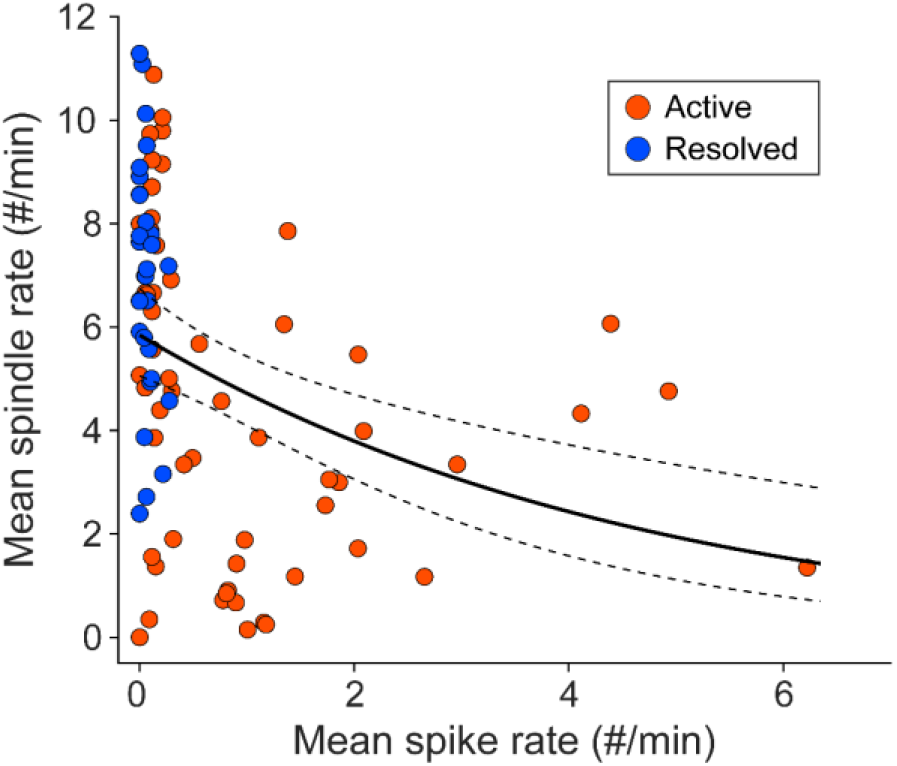
Sleep spindle rate is negatively associated with spike rate. Relationship between spindle rate and spike rate using mean rates in frontal region. Solid lines indicate age-adjusted model fits and dashed lines indicate 95% confidence intervals.

### ASM exposure is associated with higher spindle rates in active CAE and lower spindle rates in resolved CAE

In the active CAE group, untreated children had lower spindle rates than controls (p = 2.1×10^-6^, decrease of 2.5 spindles/min) and had lower spindle rates than treated children with active CAE (p = 0.04, decrease of 0.8 spindles/min; **Figure 5A**). When evaluating specific ASMs, children with active CAE treated with valproic acid (n = 7, p = 0.02, mean increase of 2.4 spindles/min) and lamotrigine (n = 6, p = 0.005, mean increase of 2.4 spindles/min; **Figure 5B**) had higher spindle rates than untreated children with active CAE. Untreated children with active CAE did not have a detectable difference in spindle rate compared to those treated with ethosuximide (n = 16, p = 0.4) or other ASMs (n = 5, p = 0.5).

**Figure 5.**
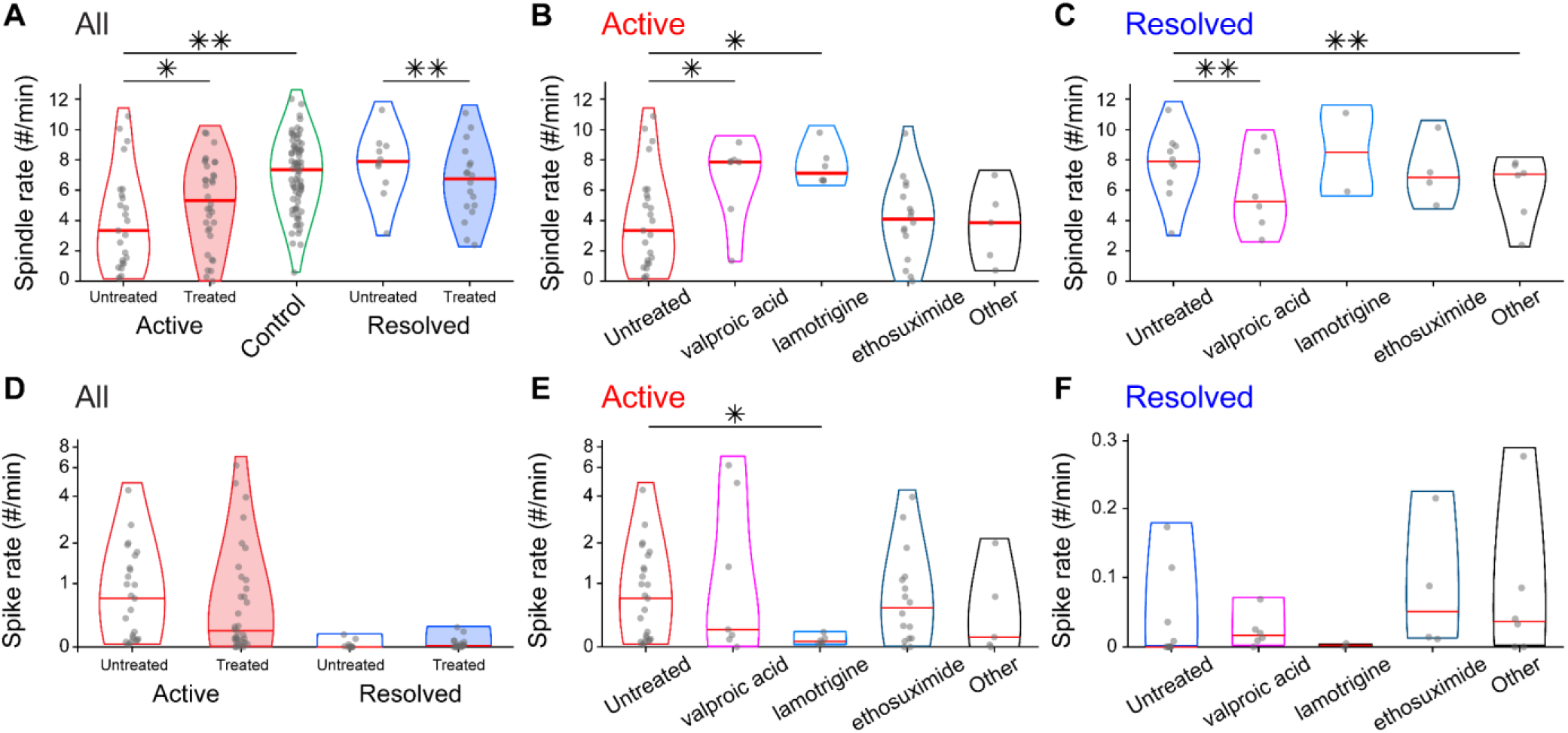
Effects of ASM on sleep spindle rate across disease states. **A)** Sleep spindle rate is lower in children with untreated active CAE than in children with treated active CAE and controls. In the resolved CAE group, sleep spindle rate is lower in treated children than in untreated children. **B)** In the active CAE group, sleep spindle rate is higher in those treated with valproic acid or lamotrigine compared with untreated children. **C)** In the resolved CAE group, sleep spindle rate is higher in untreated children than in those treated with valproic acid or other ASMs. **D)** Spike rates are shown in children with active and resolved CAE according to treatment status. **E)** In the active CAE group, spike rate is higher in children untreated than in treated children with lamotrigine. **F)** In the resolved CAE group, spike rate does not differ among medication groups. Other, Patient 1: levetiracetam; Patient 2: zonisamide; Patient 3: unknown; Patient 4: ethosuximide, valproic acid; Patient 5: lacosamide; Patient 6: ethosuximide, valproic acid. Each violin plot shows the distribution of spindle or spike rates for each group, with the red horizontal line indicating the median. *p<0.05, **p<0.001

Among children with resolved CAE spindle rates in untreated children with resolved CAE did not differ significantly from controls (p = 0.7). In contrast, treated children with resolved CAE had significantly lower spindle rates than untreated children (p = 2.2×10^-6^, decrease of 0.6 spindles/min; **Figure 5A**). When evaluating specific ASMs, children with resolved CAE treated with valproic acid (n = 6, p = 4.1×10^-26^, decrease of 0.3 spindles/min) and other ASMs (n = 6, p = 2.7×10^-17^, decrease of 0.7 spindles/min) had lower spindle rates than untreated children with resolved CAE (**Figure 5C**). No significant differences were detected between untreated resolved CAE and those treated with lamotrigine (n = 2, p = 0.5) and ethosuximide (n = 4, p = 0.5).

In contrast to spindle rate, spike rate did not differ significantly between treated and untreated children in either the active CAE group (p = 0.7) or the resolved CAE group (p = 0.5; **Figure 5D**). Among children with active CAE, those treated with lamotrigine trended to have lower spike rates than untreated children (n = 6, p = 0.04, β = -0.6), but this difference was not significant after FDR correction (FDR-adjusted p = 0.16; **Figure 5E**). No significant medication-related differences in spike rate were observed for active children treated with valproic acid (n = 7, p = 0.4), ethosuximide (n = 16, p = 0.9), or other ASMs (n = 5, p = 0.8). Among children with resolved CAE, spike rates did not differ between children treated with individual ASMs and those untreated (all p > 0.2; **Figure**

## 4. Discussion

In this study, we examined sleep spindle activity across disease states and ASM exposure in children with CAE. We found three principal results. First, children with active CAE had lower sleep spindle rates than children with resolved CAE and controls across all cortical regions, whereas spike rates were higher in children with active CAE compared to children with resolved CAE. Second, spindle rate was inversely associated with spike rate, and both spindle deficits and spike activity showed frontal predominance. Third, the relationship between ASM exposure and spindle rate differed by disease state: treated children with active CAE had higher spindle rates than untreated children, whereas treated children with resolved CAE had lower spindle rates than untreated children with resolved disease. ASM-related differences in spike rate were more limited across disease states. Together, these findings suggest that sleep spindle rate is a sensitive physiological marker of disease state in CAE and reflects both disease activity and medication-related effects.

Sleep spindles are generated through reciprocal interactions between the thalamic reticular nucleus, thalamocortical relay neurons, and cortex—the same broad circuitry implicated in generalized spike-wave discharges in CAE^20-25, 47^. This shared thalamocortical substrate has been proposed as a plausible mechanism by which epileptic activity could hijack normal sleep physiology^31^. Prior experimental and clinical studies suggest that epileptic spikes can interfere with spindle-generating networks ^25, 38^. Importantly, reduced spindle activity and spike-coupled sleep rhythms have been associated with cognitive dysfunction and impaired sleep-dependent memory consolidation^25, 26, 30, 36, 39, 48^. In contrast, spike rate alone does not reliably predict cognitive function in epilepsy^26, 30, 49-53^. More broadly, sleep spindle activity has been linked to general cognitive abilities^54^, sleep-dependent memory consolidation^27, 55^, and executive-attentional functions^56, 57^ in typically developing children. Thus, the reduced spindle activity observed in children with active CAE may contribute to the cognitive difficulties commonly reported in CAE.

Our findings of reduced spindle rate during active CAE using automated detection tools in a large cohort of children align with previous work^41, 58^ and extend this framework. Unlike the focal spindle reductions reported in regions affected by focal spikes in self-limited epilepsy with centrotemporal spikes^26, 30^, the reduction in spindle rate in CAE was observed across cortical regions, consistent with generalized discharges in CAE that broadly affect sleep architecture. Despite this widespread reduction, we also found that the most pronounced reduction in sleep spindles was observed in the frontal region, the same region with the greatest epileptic spike predominance in CAE^4, 5, 7^. Previous studies suggest that spindle activity in different cortical regions supports distinct aspects of sleep-dependent learning and memory, including contributions from frontal regions^25, 59, 60^, highlighting the importance of regional spindle activity for cognitive function^61-64^. This may be particularly relevant to CAE, as children with CAE commonly exhibit attention and executive dysfunction^9, 65^, and functional neuroimaging studies further implicate abnormalities in frontal attention networks^66^. In this study, the preferential frontal spindle deficit during active CAE may therefore be relevant to the attention and executive difficulties associated with frontal network dysfunction in CAE. In contrast, the absence of a preferential frontal spindle deficit after disease resolution suggests that these regional spindle abnormalities are state-dependent rather than a permanent feature of CAE, supporting sleep spindle activity as a dynamic marker of thalamocortical dysfunction.

ASMs, including ethosuximide, valproic acid, and lamotrigine, are generally well-tolerated treatments for CAE, with ethosuximide typically recommended as first-line therapy and valproic acid favored over lamotrigine when ethosuximide fails^11, 42, 43^. However, ASM-related effects on cognition, particularly attention, remain incompletely understood^10, 11, 43^. Randomized controlled trials have shown that ethosuximide and valproic acid are more effective than lamotrigine in controlling clinical and electrographic absence seizures, while treatment efficacy, attentional effects, and tolerability differ across medications^11, 67, 68^. Thus, although ASMs reduce electroclinical seizure burden, seizure and spike measures alone may not fully capture ASM-related differences in physiological thalamocortical activity. In the current study, treatment with valproic acid or lamotrigine was associated with higher spindle rates than no ASM treatment in active CAE. Lamotrigine was also associated with a lower spike rate. In contrast, among children with resolved CAE, valproic acid treatment was associated with lower spindle rates than no ASM treatment, indicating that valproic acid treatment may lower spindle rates after epileptiform activity has diminished. Interestingly, in the randomized controlled trial comparing valproic acid, lamotrigine and ethosuximide in children with CAE^11^, children treated with valproic acid had worse attentional outcomes than those treated with lamotrigine. Prospective studies integrating spindle measures, medication exposure, spike burden, and neuropsychological outcomes will be necessary to determine whether spindle rate can help explain medication-specific cognitive effects and guide individualized treatment decisions in CAE.

This retrospective, hospital-based study has several limitations. The sample sizes within individual ASM categories were small, limiting our power to detect medication-specific effects. Cognitive outcomes were not assessed in this study, restricting direct inference about functional consequences. Despite these limitations, the convergent findings raise the possibility that reduced spindle activity in children with active CAE may contribute to the cognitive difficulties frequently reported in this population.

In conclusion, these findings support further evaluation of sleep spindle rate as a physiological marker of thalamocortical network function that may provide information beyond epileptiform spike burden in CAE. The sensitivity of spindle activity to disease state and ASM exposure suggests that it warrants prospective evaluation as a marker of disease resolution and treatment-related changes. Prospective studies integrating spindle activity, spike burden, medication exposure, and cognitive outcomes are needed to determine its clinical relevance in CAE.

## Author Contributions

CJC planned the study; SKG and HK analyzed the data and wrote the first draft of the manuscript; SKG and CJC collected data; HK and MAK contributed to developing analysis methods; SY and DMC contributed to data analysis; HK and CJC supervised the study; all authors edited and approved the final manuscript.

## Acknowledgments

This work was supported by NINDS R01NS115868.

## Disclosure of Commercial Interests

CJC has consulted for Ovid Pharmaceuticals, Novartis, Biogen, Ionis Pharmaceuticals, and Sensorium Therapeutics and received sponsored research awards from Novartis, Biogen, Epilepsy Foundation, American Academy of Neurology, Child Neurology Foundation, Epilepsy Foundation New England, and NIH. MAK has consulted for Novartis, Biogen, Ionis Pharmaceuticals, and has received sponsored research awards from NSF, NIH, and the Burroughs Wellcome Fund.

## References

1. Specchio N, Wirrell EC, Scheffer IE, et al. International League Against Epilepsy classification and definition of epilepsy syndromes with onset in childhood: Position paper by the ILAE Task Force on Nosology and Definitions. Epilepsia 2022;63:1398–1442.

2. Loiseau P, Duché B, Pédespan J-M. Absence Epilepsies. Epilepsia 1995;36:1182–1186.

3. Grosso S, Galimberti D, Vezzosi P, et al. Childhood absence epilepsy: evolution and prognostic factors. Epilepsia 2005;46:1796–1801.

4. Elmali AD, Auvin S, Bast T, Rubboli G, Koutroumanidis M. How to diagnose and classify idiopathic (genetic) generalized epilepsies. Epileptic Disord 2020;22:399–420.

5. Hirsch E, French J, Scheffer IE, et al. ILAE definition of the Idiopathic Generalized Epilepsy Syndromes: Position statement by the ILAE Task Force on Nosology and Definitions. Epilepsia 2022;63:1475–1499.

6. Crunelli V, Leresche N. Childhood absence epilepsy: genes, channels, neurons and networks. Nature reviews Neuroscience 2002;3:371–382.

7. Sadleir LG, Farrell K, Smith S, Connolly MB, Scheffer IE. Electroclinical features of absence seizures in childhood absence epilepsy. Neurology 2006;67:413–418.

8. Crunelli V, Lőrincz ML, McCafferty C, et al. Clinical and experimental insight into pathophysiology, comorbidity and therapy of absence seizures. Brain : a journal of neurology 2020;143:2341–2368.

9. Caplan R, Siddarth P, Stahl L, et al. Childhood absence epilepsy: behavioral, cognitive, and linguistic comorbidities. Epilepsia 2008;49:1838–1846.

10. Masur D, Shinnar S, Cnaan A, et al. Pretreatment cognitive deficits and treatment effects on attention in childhood absence epilepsy. Neurology 2013;81:1572–1580.

11. Glauser TA, Cnaan A, Shinnar S, et al. Ethosuximide, valproic acid, and lamotrigine in childhood absence epilepsy. N Engl J Med 2010;362:790–799.

12. Pavone P, Bianchini R, Trifiletti RR, Incorpora G, Pavone A, Parano E. Neuropsychological assessment in children with absence epilepsy. Neurology 2001;56:1047–1051.

13. Kernan CL, Asarnow R, Siddarth P, et al. Neurocognitive profiles in children with epilepsy. Epilepsia 2012;53:2156–2163.

14. Fonseca Wald ELA, Hendriksen JGM, Drenthen GS, et al. Towards a Better Understanding of Cognitive Deficits in Absence Epilepsy: a Systematic Review and Meta-Analysis. Neuropsychology review 2019;29:421–449.

15. Shinnar RC, Shinnar S, Cnaan A, et al. Pretreatment behavior and subsequent medication effects in childhood absence epilepsy. Neurology 2017;89:1698–1706.

16. Staresina BP, Niediek J, Borger V, Surges R, Mormann F. How coupled slow oscillations, spindles and ripples coordinate neuronal processing and communication during human sleep. Nat Neurosci 2023;26:1429–1437.

17. Kwon H, Walsh KG, Berja ED, et al. Sleep spindles in the healthy brain from birth through 18 years. Sleep 2023;46.

18. Loomis AL, Harvey EN, Hobart G. POTENTIAL RHYTHMS OF THE CEREBRAL CORTEX DURING SLEEP. Science 1935;81:597–598.

19. Purcell SM, Manoach DS, Demanuele C, et al. Characterizing sleep spindles in 11,630 individuals from the National Sleep Research Resource. Nat Commun 2017;8:15930.

20. Andersen P, Andersson SA, Lomo T. Thalamo-cortical relations during spontaneous barbiturate spindles. Electroencephalogr Clin Neurophysiol 1968;24:90.

21. Buzsáki G. The thalamic clock: emergent network properties. Neuroscience 1991;41:351–364.

22. Clemente-Perez A, Makinson SR, Higashikubo B, et al. Distinct Thalamic Reticular Cell Types Differentially Modulate Normal and Pathological Cortical Rhythms. Cell Rep 2017;19:2130–2142.

23. Steriade M, McCormick DA, Sejnowski TJ. Thalamocortical oscillations in the sleeping and aroused brain. Science 1993;262:679–685.

24. Mizrahi-Kliger AD, Kaplan A, Israel Z, Bergman H. Entrainment to sleep spindles reflects dissociable patterns of connectivity between cortex and basal ganglia. Cell Rep 2022;40:111367.

25. Wodeyar A, Chinappen D, Kwon H, et al. A hierarchical cascade of sleep rhythms supports motor memory and is hijacked by epileptic spikes in human epilepsy. Proceedings of the National Academy of Sciences of the United States of America 2026;In press.

26. Kramer MA, Stoyell SM, Chinappen D, et al. Focal Sleep Spindle Deficits Reveal Focal Thalamocortical Dysfunction and Predict Cognitive Deficits in Sleep Activated Developmental Epilepsy. The Journal of neuroscience : the official journal of the Society for Neuroscience 2021;41:1816–1829.

27. Hahn MA, Heib D, Schabus M, Hoedlmoser K, Helfrich RF. Slow oscillation-spindle coupling predicts enhanced memory formation from childhood to adolescence. Elife 2020;9.

28. Clemens Z, Fabo D, Halasz P. Overnight verbal memory retention correlates with the number of sleep spindles. Neuroscience 2005;132:529–535.

29. Tamaki M, Matsuoka T, Nittono H, Hori T. Fast sleep spindle (13-15 hz) activity correlates with sleep-dependent improvement in visuomotor performance. Sleep 2008;31:204–211.

30. Kwon H, Chinappen DM, Kinard EA, et al. Association of Sleep Spindle Rate With Memory Consolidation in Children With Rolandic Epilepsy. Neurology 2025;104:e210232.

31. Beenhakker MP, Huguenard JR. Neurons that fire together also conspire together: is normal sleep circuitry hijacked to generate epilepsy? Neuron 2009;62:612–632.

32. Kostopoulos G, Gloor P, Pellegrini A, Siatitsas I. A study of the transition from spindles to spike and wave discharge in feline generalized penicillin epilepsy: EEG features. Exp Neurol 1981;73:43–54.

33. Kostopoulos G, Gloor P, Pellegrini A, Gotman J. A study of the transition from spindles to spike and wave discharge in feline generalized penicillin epilepsy: microphysiological features. Exp Neurol 1981;73:55–77.

34. Steriade M, Contreras D. Relations between cortical and thalamic cellular events during transition from sleep patterns to paroxysmal activity. The Journal of neuroscience : the official journal of the Society for Neuroscience 1995;15:623–642.

35. Frauscher B, Bernasconi N, Caldairou B, et al. Interictal Hippocampal Spiking Influences the Occurrence of Hippocampal Sleep Spindles. Sleep 2015;38:1927–1933.

36. McLaren JR, Luo Y, Kwon H, Shi W, Kramer MA, Chu CJ. Preliminary evidence of a relationship between sleep spindles and treatment response in epileptic encephalopathy. Ann Clin Transl Neurol 2023;10:1513–1524.

37. Bender AC, Jaleel A, Pellerin KR, et al. Altered Sleep Microarchitecture and Cognitive Impairment in Patients With Temporal Lobe Epilepsy. Neurology 2023;101:e2376–e2387.

38. Wodeyar A, Chinappen D, Mylonas D, et al. Thalamic epileptic spikes disrupt sleep spindles in patients with epileptic encephalopathy. Brain : a journal of neurology 2024;147:2803–2816.

39. Spencer ER, Chinappen D, Emerton BC, et al. Source EEG reveals that Rolandic epilepsy is a regional epileptic encephalopathy. NeuroImage Clinical 2022;33:102956.

40. Stoyell SM, Baxter BS, McLaren J, et al. Diazepam induced sleep spindle increase correlates with cognitive recovery in a child with epileptic encephalopathy. BMC Neurology 2021;21:355.

41. Zhang W, Xin M, Song G, Liang J. Childhood absence epilepsy patients with cognitive impairment have decreased sleep spindle density. Sleep Medicine 2023;103:89–97.

42. Brigo F, Igwe SC, Lattanzi S. Ethosuximide, sodium valproate or lamotrigine for absence seizures in children and adolescents. Cochrane Database Syst Rev 2021;1:Cd003032.

43. Kessler SK, McGinnis E. A Practical Guide to Treatment of Childhood Absence Epilepsy. Paediatr Drugs 2019;21:15–24.

44. Wirrell EC, Camfield CS, Camfield PR, Gordon KE, Dooley JM. Long-term prognosis of typical childhood absence epilepsy. Neurology 1996;47:912–918.

45. Helfrich RF, Lendner JD, Mander BA, et al. Bidirectional prefrontal-hippocampal dynamics organize information transfer during sleep in humans. Nat Commun 2019;10:3572.

46. Conrad EC, Revell AY, Greenblatt AS, et al. Spike patterns surrounding sleep and seizures localize the seizure-onset zone in focal epilepsy. Epilepsia 2023;64:754–768.

47. Merricks EM, Kwon H, Wang P, Walsh KG, Kramer MA, Chu CJ. Corticothalamic loops and cellular networks: implications for thalamic neuromodulation in epilepsy. Neurotherapeutics 2026.

48. Kwon H, Merricks EM, Yi S, Kramer MA, Chu JC. Sleep microarchitecture, epileptic spikes, and memory in epilepsy: a unifying framework with implications for developmental and epileptic encephalopathies. Journal of Clinical Neurophysiology 2026.

49. Yi JD, Pasdarnavab M, Kueck L, Tarcsay G, Ewell LA. Behavioral Timing of Interictal Spikes, But Not Rate, Correlates with Impaired Working Memory Performance. The Journal of neuroscience : the official journal of the Society for Neuroscience 2025;45.

50. Ebus S, Arends J, Hendriksen J, et al. Cognitive effects of interictal epileptiform discharges in children. Eur J Paediatr Neurol 2012;16:697–706.

51. Bjornaes H, Bakke KA, Larsson PG, et al. Subclinical epileptiform activity in children with electrical status epilepticus during sleep: effects on cognition and behavior before and after treatment with levetiracetam. Epilepsy Behav 2013;27:40–48.

52. van Arnhem MML, van den Munckhof B, Arzimanoglou A, et al. Corticosteroids versus clobazam for treatment of children with epileptic encephalopathy with spike-wave activation in sleep (RESCUE ESES): a multicentre randomised controlled trial. The Lancet Neurology 2024;23:147–156.

53. Vega C, Sanchez Fernandez I, Peters J, et al. Response to clobazam in continuous spike-wave during sleep. Dev Med Child Neurol 2018;60:283–289.

54. Hoedlmoser K, Heib DP, Roell J, et al. Slow sleep spindle activity, declarative memory, and general cognitive abilities in children. Sleep 2014;37:1501–1512.

55. Astill RG, Piantoni G, Raymann RJ, et al. Sleep spindle and slow wave frequency reflect motor skill performance in primary school-age children. Frontiers in human neuroscience 2014;8:910.

56. Morales-Ghinaglia M, He F, Calhoun SL, et al. Association of sleep spindle activity with executive functioning and intellectual ability in children and adolescents. Sleep 2025;48.

57. Saito Y, Kaga Y, Nakagawa E, et al. Association of inattention with slow-spindle density in sleep EEG of children with attention deficit-hyperactivity disorder. Brain Dev 2019;41:751–759.

58. Myatchin I, Lagae L. Sleep spindle abnormalities in children with generalized spike-wave discharges. Pediatr Neurol 2007;36:106–111.

59. Born J, Wilhelm I. System consolidation of memory during sleep. Psychol Res 2012;76:192–203.

60. Staresina BP. Coupled sleep rhythms for memory consolidation. Trends in cognitive sciences 2024;28:339–351.

61. Schmidt C, Peigneux P, Muto V, et al. Encoding difficulty promotes postlearning changes in sleep spindle activity during napping. The Journal of neuroscience : the official journal of the Society for Neuroscience 2006;26:8976–8982.

62. Gais S, Molle M, Helms K, Born J. Learning-dependent increases in sleep spindle density. The Journal of neuroscience : the official journal of the Society for Neuroscience 2002;22:6830–6834.

63. Fernandez LMJ, Lüthi A. Sleep Spindles: Mechanisms and Functions. Physiological Reviews 2020;100:805–868.

64. Morin A, Doyon J, Dostie V, et al. Motor sequence learning increases sleep spindles and fast frequencies in post-training sleep. Sleep 2008;31:1149–1156.

65. Dlugos D, Shinnar S, Cnaan A, et al. Pretreatment EEG in childhood absence epilepsy: associations with attention and treatment outcome. Neurology 2013;81:150–156.

66. Killory BD, Bai X, Negishi M, et al. Impaired attention and network connectivity in childhood absence epilepsy. Neuroimage 2011;56:2209–2217.

67. Cnaan A, Shinnar S, Arya R, et al. Second monotherapy in childhood absence epilepsy. Neurology 2017;88:182–190.

68. Glauser TA, Cnaan A, Shinnar S, et al. Ethosuximide, valproic acid, and lamotrigine in childhood absence epilepsy: initial monotherapy outcomes at 12 months. Epilepsia 2013;54:141–155.

